# Observing Mechanosensitive Channels in Action in Living Bacteria

**DOI:** 10.1101/2023.08.24.554739

**Authors:** Mohammad Sharifian Gh., Michael J. Wilhelm, Hai-Lung Dai

## Abstract

Mechanosensitive (MS) channels act to protect the cytoplasmic membrane (CM) of living cells from environmental changes in osmolarity. In this report, we demonstrate the use of time-resolved second-harmonic light scattering (SHS) as a means of experimentally observing the relative state (open vs. closed) of MS channels in living bacteria suspended in different buffer solutions. Specifically, the state of the MS channels was selectively controlled by changing the composition of the suspension medium, inducing either a transient or persistent osmotic shock. SHS was then used to monitor transport of the SHG-active cation, malachite green (MG) across the bacterial CM. When MS channels were forced open, MG cations were able to cross the CM at a rate at least two orders of magnitude faster compared to when the MS channels were closed. These observations were corroborated using both numerical model simulations and complementary fluorescence experiments, in which the propensity for the CM impermeant cation, propidium to stain cells was shown to be contingent upon the relative state of the MS channels (i.e., cells with open MS channels fluoresced red, cells with closed MS channels did not). Application of time-resolved SHS to experimentally distinguish MS channels opened via osmotic shock vs. chemical activation, as well as a general comparison to the patch-clamp method is discussed.

## INTRODUCTION

Mechanosensitive (MS) channels^1^ are membrane-gated proteins responsible for preserving the integrity of the cytoplasmic membrane (CM) in response to environmental shifts in osmolarity in a variety of cell types.^2–4^ In eukaryotes, MS channels are involved in diverse processes such as embryonic development, touch, pain, hearing, lung growth, and muscle homeostasis.^5–8^ In bacteria, MS channels are fundamental components of the CM that play a critical role as a ‘safety valve’ under hypo-osmotic shock (osmotic downshock).^1,9,10^ They respond to acute changes in lateral tension in the membrane lipid bilayer generated by rapid diffusion of water into the cytosol and create transient pores to release the pressure by rapid equilibration of internal osmolytes (e.g., potassium and glutamate)^11–13^ and external solutes (e.g., sodium and hydrogen ions).^14,15^ Bacterial MS channels are primarily categorized by function into two major families, the MS channels of small conductance, MscS (1 nanosiemens conductance; 1.8 nm pore diameter)^16–19^ and the MS channels of large conductance, MscL (3.5 nanosiemens conductance; 3 nm pore diameter).^15,20–22^ The channels are generally non-specific in terms of the ions and molecules that pass through the open pores.^9^ Once in the open state, the MS channel permits passage of any molecule sufficiently small to pass through. For the CM of *E. coli* for example, it has been reported that there are roughly 1200 MS channels (about 560 MscL and 610 MscS channels) within a single bacterium.^23,24^

According to the van’t Hoff Law, the osmotic pressure ∏ can be expressed as *RT* ∑ *C*_*i*_, where *R* is the universal gas constant, *T* is the absolute temperature, and *C*_*i*_ is the molar concentration of solute *i* (i.e., the osmolarity, Osm). Thus, the osmotic pressure sensed by a biological membrane stems from the difference in the osmolarity, Δ∏ = |*∏*_*out*_ *- ∏*_*ln*_ | across the membrane.^25,26^ For example, a mere 40 mOsm difference across a cell membrane results in a pressure of roughly 1 atm. At this pressure, without a suitable mechanism to retain the structural integrity of the cell membrane during the extreme increase in cell turgor, the cell would burst within seconds.^9,27–29^ As depicted in **Scheme 1**, in response to such changes in pressure, MS channels are automatically pulled open, allowing the release of intracellular solute molecules and hence re-establishment of osmotic equilibrium across the membrane.

In addition to several *in silico* studies,^30,31^ various experimental approaches, including patch-clamp electrophysiology,^4,32–34^ single molecule Förster resonance energy transfer (FRET),^35,36^ and electron paramagnetic resonance (EPR) spectroscopy^3,20,37^ have been employed to study the kinetics and thermodynamics of MS channels. The patch-clamp technique has been widely used to measure the conductance, open dwell time, pressure sensitivity, and ion selectivity of MS channels by employing simplified membrane systems, including: giant bacterial protoplasts^1^ and spheroplasts,^38,39^ reconstituted channels in giant unilamellar liposomes,^40,41^ and membranes fused with liposomes.^42^ In this experimental approach, the pressure applied to the pipette tip can be controlled precisely but how it is converted to local stress varies in each experiment. Indeed, the assumption for interpreting data with this approach is that MS channels respond to membrane tension, not to hydrostatic pressure perpendicular to it.^43,44^ In fluorescence-based techniques, the incorporation of a fluorescent label may induce nontrivial changes in the system. Further, EPR-based structural analysis is limited to reconstituted proteins in liposomes.

Currently, there are no electrophysiological methods capable of studying intact bacteria in a non-destructive manner during downshock. In this report, we introduce a nonlinear optical technique which is capable of monitoring MS channels in action in live bacteria without incorporation of a fluorescent label and/or non-osmotic pressure. Specifically, second-harmonic light scattering (SHS), a surface-sensitive technique, has been demonstrated to be able to monitor in real time molecular adsorption and transport across membranes in living cells.^45–55^ As conditions are employed to create an osmotic down shock to the membranes of a living bacterium, SHS can be used to observe transport of molecules across the MS channels.

**Scheme 1.**
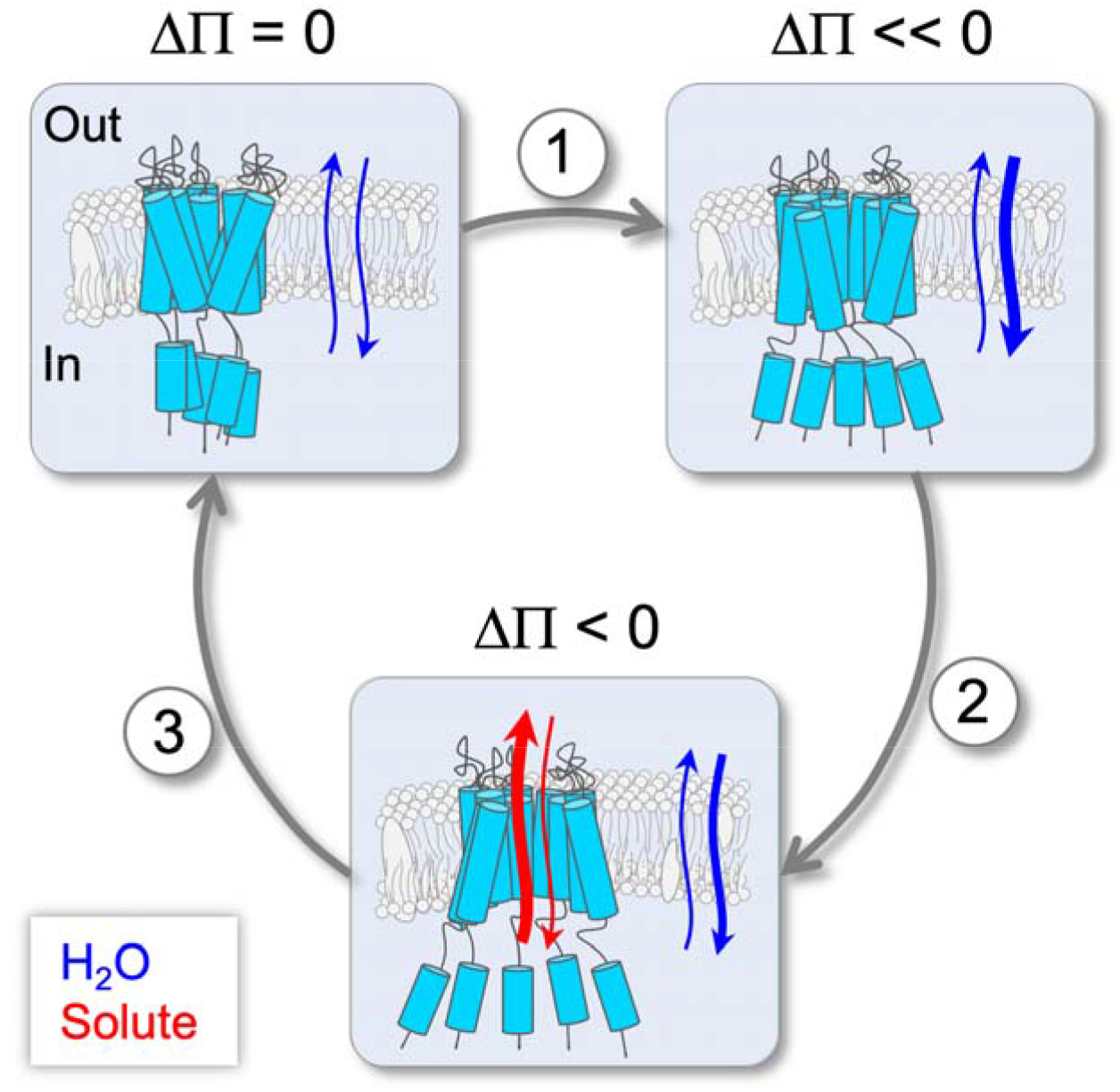
Cartoon schematic depicting osmotic shock-induced activation (Δ∏<<0) of MS channels, along with the associated changes in the diffusion of water (blue arrows) and solute (red arrows) across the cytoplasmic membrane and MS channels, respectively.

The phenomenon of and theoretical basis for SHS has already been described in detail.^56–60^ Briefly, SHS is based on the nonlinear optical phenomenon, second-harmonic generation (SHG) in which a fraction of an incident light of frequency is scattered at frequency 2 after interacting with second-harmonic (SH) active matter (see **Scheme 2A**), such as molecules that lack center-of-inversion symmetry (e.g., the malachite green cation, MG). **Scheme 2B** shows the characteristic time-resolved SHS signal for the molecular transport response of an SH-active molecule passively diffusing across a cell membrane. Of significance, an ensemble of SH-active molecules isotropically oriented in solution produces no SH signal: while each molecule may produce SH light, the SH optical fields produced by oppositely oriented nearest neighbor molecules result in destructive interference. Nevertheless, alignment of the molecules on the outer leaflet of the membrane results in detectable SH signal due to constructive interference of SH fileds.^56–60^ Further, if the molecule is membrane-permeable and able to adsorb on the inner leaflet of the membrane, SH fields generated from oppositely oriented molecules on the two leaflets will destructively interfere, resulting in an attenuation of the SH signal over the course of transport. This sequential rise and decay of signal is the characteristic SH response for surface adsorption and transport of the SH-active molecule across the cell membrane. This interpretation has been validated in numerous SHS-based studies of liposomes^61–66^ and living cells,^45,46,55,67–70,47–54^ as well as confirmed with brightfield transmission microscopy.^49^

**Scheme 2.**
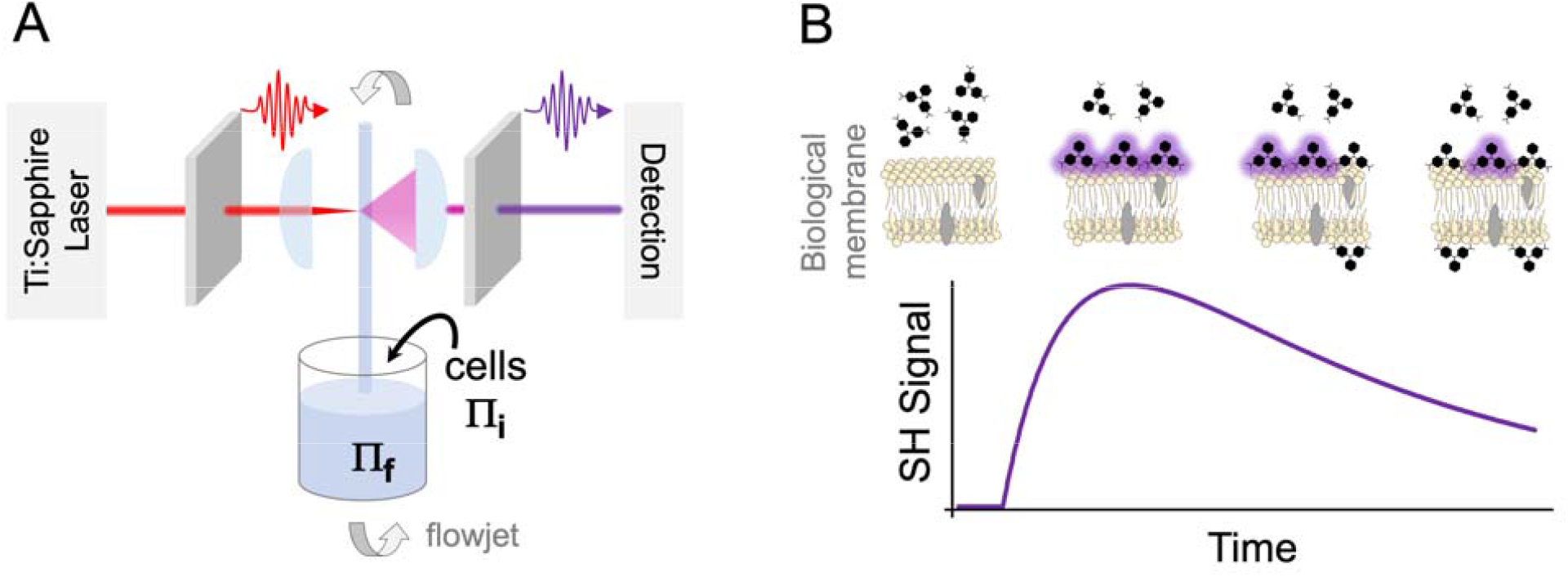
(A) Schematic of the time-resolved SHS experiment and (B) representative time-resolved SHS response of SHG-active molecules interacting with the opposing leaflets of a biological membrane.

Herein, we show that time-resolved SHS can be used to experimentally observe the state (i.e., open *vs*. closed) of bacterial MS channels in a colloidal suspension of living bacteria. Our lab has previously demonstrated that perturbations to membrane permeability, whether from antimicrobial attack^51,52^ or activation of membrane embedded proteins,^47^ can be quantitatively deduced from characteristic changes observed in the molecular transport kinetics. Likewise, for observing the mechanosensitive channels in action, we use an SHG-active probe molecule (i.e., the MG cation) which is sufficiently small (ca. 1 nm wide) and capable of crossing through an open MS channel of *E. coli*. When the MS channel is activated, it provides an efficient alternative route for diffusive transport across the bacterial CM and should therefore result in characteristic perturbations in the SHS kinetic response. Application of time-resolved SHS as a new experimental paradigm for examination of MS channels and MS channel activators in living microorganisms is demonstrated.

## MATERIALS and METHODS

### A. Sample preparation

A sterile loop of wild-type *E. coli* strain, AW405^71^ was used to streak a Luria Broth agar plate (Sigma-Aldrich, L2897) which was then allowed to aerobically incubate at 37°C for 24 hours to enable colonization. For each SHS experiment, a discrete colony of the bacterial strain was cultivated aerobically at 37°C in 50 ml of Terrific Broth, TB culture medium (Sigma-Aldrich, T0918), in a shaking flask (150 rpm) for 8 hours. This growth period corresponds to the middle-to-late exponential phase. Cells underwent a gentle centrifugation process (1500×g, 2 minutes, room temperature) and pellets were subjected to two rounds of washing with an appropriate volume of phosphate-buffered saline (1×PBS; pH 7.3) to eliminate waste and residual TB. During each washing step, a Rotamix device (Model 10101-RKVSD, ATR Inc.) operating at 20 rpm was utilized to resuspend the bacterial cells in 1×PBS, ensuring that no biomechanical forces were exerted on the bacteria during the resuspension process. After thorough washing with 1×PBS, the supernatant was discarded, and the resulting pellets were collected for the preparation of *E. coli* samples in 1×PBS. The optical density (at 600 nm light wavelength) of the bacterial suspension was measured to ascertain the suitable final densities for each SHS experiment.

Stock solutions of malachite green oxalate (Sigma Aldrich, M9015) were prepared using distilled deionized water (Millipore, 18.2 MΩ·cm) and stored in a dark environment at 4°C for use.

### B. Time-resolved SHS

The setup for the SHS experiments has been described in detail previously.^49^ Briefly, the fundamental light source employed was the 800 nm pulses (pulse width 150 fs, repetition rate 76 MHz, pulse energy 5 nJ) from a mode-locked Ti:Sapphire laser (Coherent, Micra V, oscillator only). To collect the SH signal while suppressing all other scattered light background, a BG39 band-pass filter and a monochromator with a bandwidth of 400±1 nm (1 mm entrance and exit slits) were utilized. Additionally, to distinguish the coherent SHG response from the hyper-Rayleigh background scattering (HRS) originating from the bulk dye solution, an initial measurement was performed solely on the MG solution without any bacteria cell added. Signal measurements were taken at regular intervals of 1.54 seconds, with a gate time of 1.0 second.

To minimize absorption losses of the laser and the signal and to avoid SH scattering which often occurs at interfaces, a liquid jet column produced by a liquid flow system, instead of a liquid in a cuvette, was used as samples. This liquid jet column was generated by pumping the sample solution through a circular stainless-steel nozzle with an inner diameter of 1.59 mm. Nalgene tubing (Nalge Nunc, Inc.) was used to establish connections between the sample reservoir and the inlet of a motorized liquid pump (Micropump, Inc.), as well as for recollecting the sample back into the reservoir (see **Scheme 2A**). A magnetic stirrer (Spectrocell, Inc.) was employed to ensure efficient mixing within the sample reservoir.

Prior to each SHS experiment, an appropriate amount of the stock solution was utilized to achieve the desired concentration of MG in the liquid flow jet. A small quantity of bacteria stock suspension was then introduced into the flowing MG solution at t=0 s to achieve a bacteria cell density of (2.5 ± 0.1)×10^8^ cells.ml^-1^. All experiments were conducted at a room temperature of 295K.

### C. Brightfield and fluorescence imaging

In each imaging experiment, bacterial suspensions under different conditions were subjected to incubation with a membrane-impermeable fluorescence marker, propidium iodide (Pro) at a concentration of 20 μM (Sigma-Aldrich, P4170). The incubation process was carried out in the dark at room temperature for a duration of 15 minutes. Subsequently, a 20 μl aliquot of each sample was placed on a microscope glass slide, covered with a glass coverslip, and mounted onto the fluorescence microscope stage. For each glass slide (representing a sample), epi-fluorescence images were captured from a minimum of 15 field-of-view (FOV). In each experiment, a substantial number of cells were enumerated. The Pro-stained bacteria observed in the images indicate the presence of membrane permeabilized bacteria. The imaging procedure utilized a Leica DMRXE microscope operating in brightfield transmission mode. A 100× PlanApo objective lens was employed in conjunction with a digital image capture system (Tucsen model TC-3) which was controlled by TSView software (OnFocus Laboratories, ver. 7). Excitation of the Pro was accomplished using an EXFO X-cite 120 Fluorescence Illuminator light source, and the resulting red fluorescence emissions were recorded in an epi-fluorescence configuration using appropriate filter cubes. The employed filter cubes possessed specific excitation and detection wavelengths centered at 560 and 630 nm, respectively (CHROMA, 49008). Image analysis was conducted using ImageJ software (version 1.43u), developed by the National Institutes of Health. The red color was changed to white in order to enhance its compatibility with the black color background, thereby improving the overall intuitiveness.

## RESULTS and ANALYSIS

In order to selectively observe molecular transport across bacterial MS channels, we consider three specific experimental scenarios for our SHS experiments: 1) bacterial suspensions in which the MS channels are closed, 2) bacterial suspensions in which the MS channels are held open, and 3) bacterial suspensions in which the MS channels were held open for a prolonged duration (ca. 30 min) and then allowed to close. It is important to note, for all three scenarios considered, all of the bacteria experience some degree of osmotic shock. The critical differences are the duration of the applied shock and whether the MS channels are open while MG cations are diffusing across the CM during the SHS experiment.

In the first scenario (i.e., closed MS channels), the bacteria experience only a transient osmotic shock as the stock sample of cells suspended in 1xPBS were added to a solution of MG cations dissolved in distilled water. Under these conditions, the MS channels briefly open and then rapidly close once equilibrium of osmolytes across the CM was re-established.^72^ This scenario represents the typical experimental conditions experienced by the bacteria in all of our previously reported SHS experiments.^45,46,55,47–54^ Importantly, despite the fact that the MS channels were briefly opened, by the time MG began to diffuse across the CM, the MS channels were expected to be closed.

In the second scenario (i.e., open MS channels), the bacteria experienced a prolonged osmotic shock. Rather than 1xPBS, the stock sample of cells were suspended in distilled water (for 30 min) and then an aliquot was added to a solution of MG cations in distilled water. Under these conditions, the MS channels were forced open and could not close. As distinct from the first scenario, when MG cations arrived at the CM, the MS channels would already be open and provide an alternative (and presumably faster) route to cross into the cytosol.

In the third and final scenario (i.e., MS channels held open and then allowed to close), the bacteria first experienced a prolonged osmotic shock, identical to the conditions of the second scenario. However, before the start of the SHS experiment, the cells were spun down into a pellet and re-suspended in 1xPBS, similar to the cells in the first scenario. An aliquot of this stock sample was then added to a solution of MG cations in distilled water. Similar to the first scenario, these bacteria experienced a transient osmotic shock at the beginning of the SHS experiment. However, by the time MG cations began to cross the CM, the MS channels were once again anticipated to be closed.

The relative state of the MS channels (i.e., open vs. closed) is expected to have a substantial influence on the observed transport kinetics across the bacterial CM. Specifically, when the MS channels are closed, as in the case of the first and third scenarios, it is postulated that MG cations will only slowly diffuse across the hydrophobic interior of the bacterial CM (i.e., similar to what has previously been reported in all of our prior studies). Conversely, when the MS channels are open, it is predicted that the MG cations will rapidly diffuse across the CM through the open MS channels, which will be reflected in the SHS signal as an attenuation of the CM transport peak (the extent of which will be proportional to the rate at which the MG cations cross the MS channels).

### A. MS Channels Open in Response to Osmotic Shock

**Figure 1** illustrates representative time-resolved SHS kinetic responses observed for MG cations interacting with *E. coli* under the three distinct scenarios detailed above. Specifically, the red trace corresponds to scenario one (transient osmotic shock, MS channels closed), the blue trace scenario two (persistent osmotic shock, MS channels open), and the green trace scenario three (persistent osmotic shock followed by relaxation, closed MS channels).

**Figure 1.**
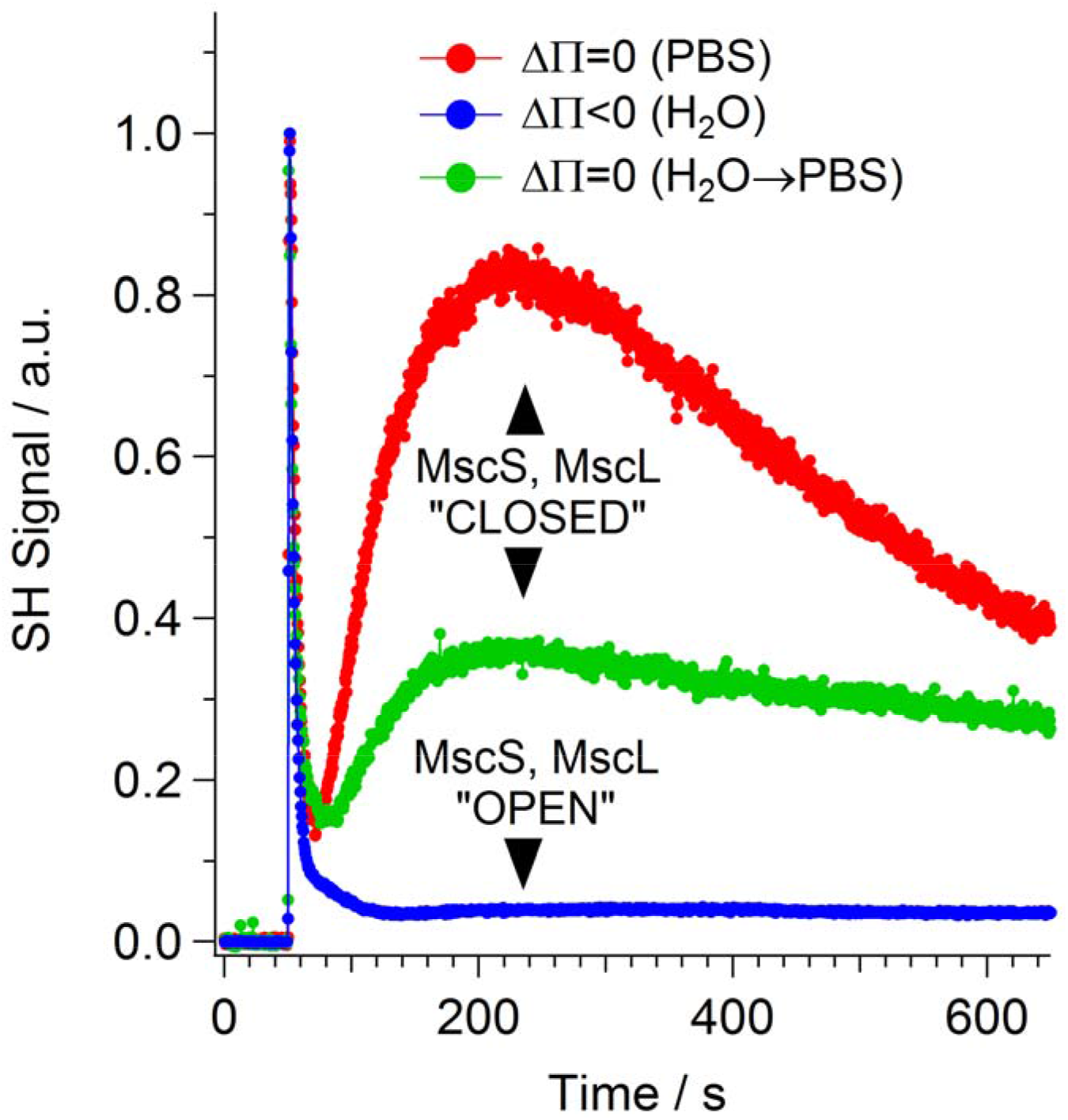
Measured time-resolved SHS traces for MG interacting with E. coli experiencing different osmolarity changes including, transient osmotic shock (red curve), persistent osmotic shock (blue curve), and persistent shock followed by cessation of shock (green curve).

Consistent with our prior SHS reports,^45–55^ the red trace corresponds to the interaction of MG cations with *E. coli* experiencing only a transient osmotic shock. Importantly, on the time scale of the SHS experiment, the MS channels are anticipated to be closed. As shown in **Figure 1**, the measured SHS signal consists of two transport events (i.e., rise and decay of signal), which can be assigned to passive diffusion across the bacterial OM and CM. Specifically, the cells were added to the system near t=50 seconds, yielding an immediate rise of SHS signal as MG cations saturated the exterior surface of the OM. This is followed by an equally rapid decay of signal (over roughly 20 seconds) as MG cations crossed the outer membrane protein (Omp) channels and adsorbed onto the interior leaflet of the OM, resulting in coherent cancellation of the SHS response. Thereafter, the SHS signal exhibits a subsequent slower secondary rise from ca. t = 70 s to t = 200 s, followed by an even more gradual decay which continues for the remainder of the observation. The secondary rise is assignable to slow diffusion across the bacterial peptidoglycan mesh (PM) and adsorption onto the exterior surface of the CM. Likewise, the slow secondary decay corresponds to the gradual diffusion of MG across the hydrophobic interior of the CM and the eventual adsorption of MG cations onto the interior leaflet of the CM. The observed slow CM transport response suggests that the MS channels have closed following the transient osmotic shock and hence do not contribute to the transport of MG ions across the CM.

Conversely, the blue trace in **Figure 1** corresponds to scenario two, in which the bacteria experienced persistent osmotic shock and hence the MS channels were held open during the full course of the experiment. Similar to the transient osmotic shock case (red trace), the SHS response for MG cations interacting with cells experiencing persistent osmotic shock begins with a rapid transport event, assignable to MG cations diffusing across the OM. The initial rise and decay of the signal is virtually identical to what was observed for the transient osmotic shock case (**Figure 1**, red trace). This is reasonable given that the MS channels are localized to the CM and hence their relative state should have no influence on molecular transport across the OM. Following the decay of the OM transport peak, the measured SHS signal simply exhibits a stable baseline plateau.

In order to determine whether the distinct SHS responses shown in **Figure 1** can be attributed to the activity of the relative state of the MS channels (i.e., open vs. closed), a series of simulated SHS responses were calculated. **Figure 2** depicts a series of simulated SHS kinetic responses corresponding to transport of MG cations across the bacterial OM, PM, and CM in a typical Gram-negative bacterium. The simulations were calculated using our previously described kinetic model of molecular uptake in Gram-negative bacteria, in which the model parameters were set according to prior experimental measurements.^45–55^ The adsorption and transport rates employed in the simulations (**Table 1**) are typical of what has previously been measured for MG interacting with *E. coli* and were held constant across all simulations.^45–55^ To exemplify the effect of the MS channels, in addition to passive diffusion across the CM, the simulations also incorporated a rate process for direct transport of MG cations across MS channels embedded in the CM. The associated transport rate across the MS channels (i.e., k_ms_) was selectively varied in each of the simulations from k_ms_ = 0 corresponding to MS channels closed (red trace) up to a maximum of k_ms_ = 0.08 s^-1^ for transport through open MS channels (purple trace).

**Table 1.** Transport rates used to simulate SHS signals. With the exception of the transport rate across the MS channels (k_ms_), all other rates were held constant across all simulations.

| Transport Medium | Term | Transport Rate ($\text{s}^{-1}$) |
| --- | --- | --- |
| Omp Channels | $k_{\text{Omp}}$ | $7 \times 10^{-2}$ |
| Peptidoglycan Mesh | $k_{\text{PM}}$ | $5 \times 10^{-2}$ |
| Cytoplasmic Membrane | $k_{\text{CM}}$ | $2 \times 10^{-4}$ |
| MS Channels | $k_{ms}$ | (0, $4 \times 10^{-4}$, $4 \times 10^{-3}$, $4 \times 10^{-2}$, $8 \times 10^{-2}$) |

**Figure 2.**
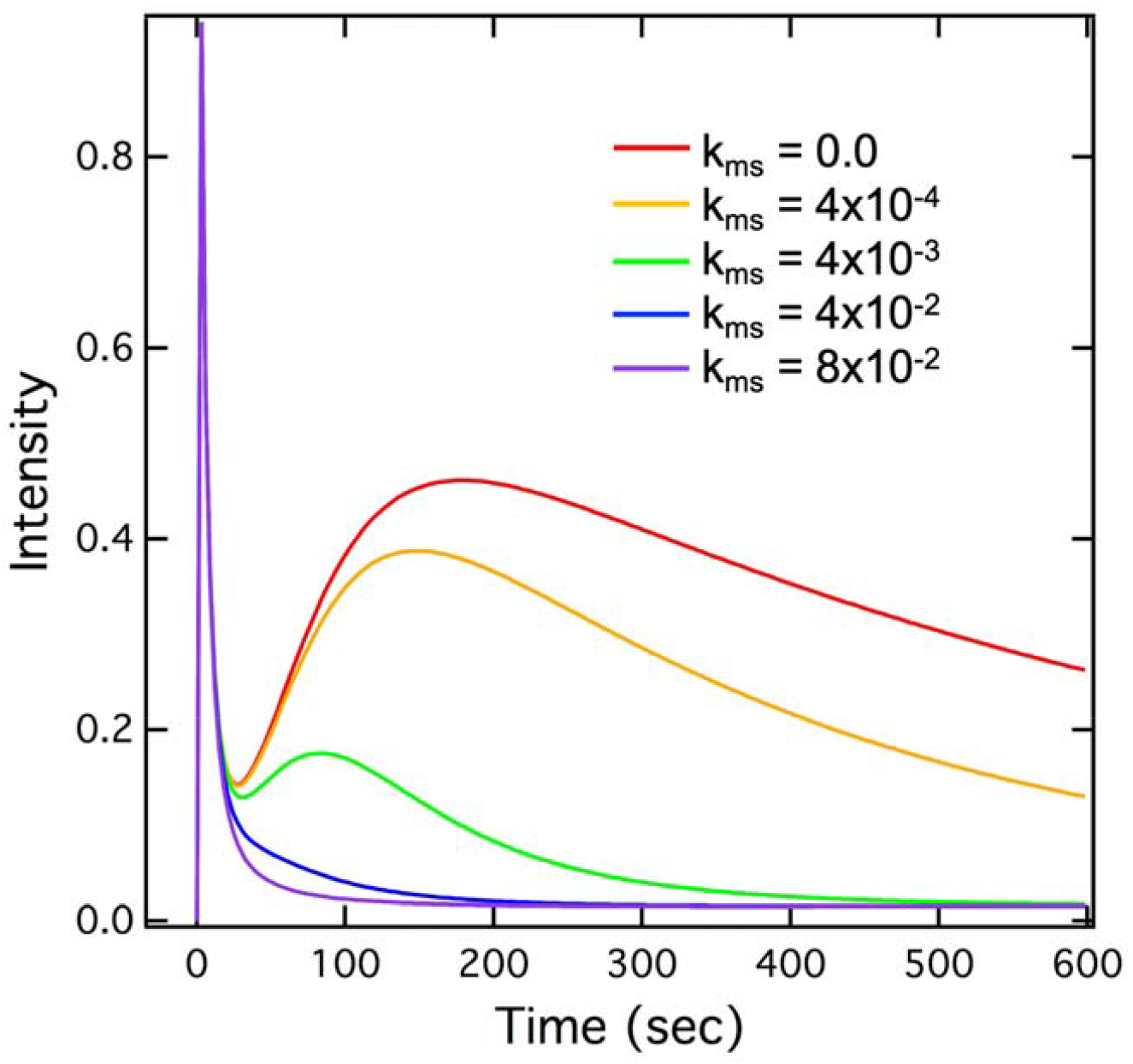
Simulated SHS traces, based upon a model for Gram-negative bacteria molecular uptake, for which the transport rate across MS channels (k_ms_) is gradually increased.

Given that MS channels are localized to the CM, variation of k_ms_ had no influence on the OM transport peak in the simulations. Conversely, as k_ms_ was increased, the SHS transport response across the CM was observed to decay at a faster rate and the magnitude of the predicted CM transport peak was shown to decrease. Of particular interest, when k_ms_ was comparable in magnitude to the typical transport rate for MG crossing the topologically similar Omp channels (ca. 0.07 s^-1^), the CM transport peak was completely lost (purple trace). Physically, the disappearance of the CM transport peak at faster MS transport rates is a result of simultaneous saturation of MG cations on both the interior and exterior leaflets of the CM. Under these conditions, there is no net abundance of MG adsorbed on either side of the membrane and hence the SHS signal is always coherently cancelled. It is important to note that the absence of the second rise-decay in the SHS trace does not suggest that MG is not transporting across the MS. Rather, these simulations predict that if MG is able to cross the MS at a sufficiently rapid rate, the CM transport peak will simply not be observed in the resulting SHS kinetic response.

The predictions of the simulations (**Figure 2**) are fully consistent with what was observed in the measured SHS kinetic responses (**Figure 1**) for MG interacting with *E. coli* prepared under scenarios one (MS channels closed, red trace) and two (MS channels open, blue trace). In particular, the fact that the CM transport peak is completely lost in scenario two (MS channels open) is consistent with the condition that the MS channels are open and that MG cations are rapidly diffusing across them. Moreover, based upon the simulated rates, the observed loss of the CM transport peak suggests that MG cations transport across the open MS channels with a rate at least as fast as that observed for the Omp channels in the OM (ca. ≥0.07 s^-1^). It is important to stress that this represents a two-orders-of-magnitude increase in the total transport rate of MG cations across the CM. Furthermore, this rate for crossing the open MS channels necessarily corresponds to a lower bound estimate, as faster transport rates would yield indistinguishable results. Nevertheless, given the similar topologies of the open MS channels in the CM and Omp channels in the OM (i.e., pore diameters of ca. 2+ nm),^73,74^ it is reasonable that MG cations should exhibit similar transport rates for crossing both of these non-selective channels.

### B. Reversibility of Prolonged Osmotic Shock

A concern over exposing bacteria to conditions of persistent osmotic shock (e.g., by suspending them in distilled deionized water) is whether their CM will simply burst. If so, the absence of the CM transport signal observed in scenario two (**Figure 1**, blue trace) could be a result of the obliteration of the bacterial CM. The third experimental scenario was designed to test this possibility. As detailed above, scenario three consists of cells exposed to a prolonged osmotic shock (i.e., identical to what was experienced by the cells in scenario two) but were then resuspended in 1xPBS (i.e., similar to the cells in scenario one), in which their MS channels were allowed to close. Scenario three also tests whether the effects of persistent osmotic shock (i.e., scenario two) are reversible.

If persistent osmotic shock had indeed caused the bacterial CM to burst, this would be an irreversible process and should show no change following re-suspension in 1xPBS. Conversely, if persistent osmotic shock simply held the MS channels open, the CM would remain intact and MG transport across the hydrophobic interior of the CM should be observable. Indeed, as shown in **Figure 1** (green trace), when cells exposed to prolonged osmotic shock are allowed to relax (scenario three, closed MS channels), the CM transport peak is once again observed. This suggests that the bacterial CM largely maintained its structural integrity during the application of the prolonged osmotic shock. It is of interest to note, however, that the magnitude of the CM peak is not as great as was observed for the cells in scenario one (red trace, transient osmotic shock). Additionally, the decay of the CM transport peak is notably slower compared to that of the cells in scenario one. These perturbations in the measured kinetic responses likely stem from residual damage to the CM following prolonged osmotic shock and will be discussed below.

Finally, as a test of the viability and morphology of these cells, we also ran complementary brightfield transmission and propidium (Pro) stained fluorescence images of the three distinct sample cases. As shown in **Figure 3**, regardless of the composition of the suspension medium, the bacterial cells all appear to exhibit similar morphology. In particular we note that the cells exposed to persistent osmotic shock (scenario two, MS channels open) are clearly still intact. However, it is possible that, because the OM and PM remain reasonably undamaged, such images are unable to definitively characterize the structural integrity of the CM. Nevertheless, if the MS channels are held in a persistent open state, it is reasonable to speculate that the CM impermeant ion, Pro, should likewise be able to cross the bacterial CM (regardless of the viability of the cell). Indeed, as shown in **Figure 3**, virtually all of the cells exposed to persistent shock (scenario two, open MS channels) fluoresced red. Conversely, only a portion (ca. 20%) of the cells experiencing transient osmotic shock (scenario one, closed MS channels) fluoresced red. This was likewise observed for the cells that experienced prolonged shock but were then allowed to relax (scenario three, closed MS channels), only a small portion fluoresced red. Consistent with the interpretation of the SHS results of MG cations shown in **Figure 1**, the Pro results suggest that Pro cations were also able to cross the CM in living cells through their open MS channels and hence fluorescence red. Specifically, when the MS channels were held open (scenario two, persistent osmotic shock), all of the cells fluoresced red. However, by closing the MS channels (by resuspending the cells in PBS, scenario three), Pro was no longer able to cross the CM and hence the majority of the cells (ca. 80%) did not fluoresce red. Importantly, this test demonstrates reversibility of the propensity for Pro to cross the bacterial CM, based solely upon the composition of the suspension medium. This interpretation suggests that suspending the bacteria in distilled deionized water held them in a state of persistent osmotic shock, in which their MS channels were forcibly held open and hence permitted free diffusion of otherwise CM impermeable molecules. However, by spinning the cells down into a pellet and resuspending them in 1xPBS, the cells were able to relax and their MS channels closed.

**Figure 3.**
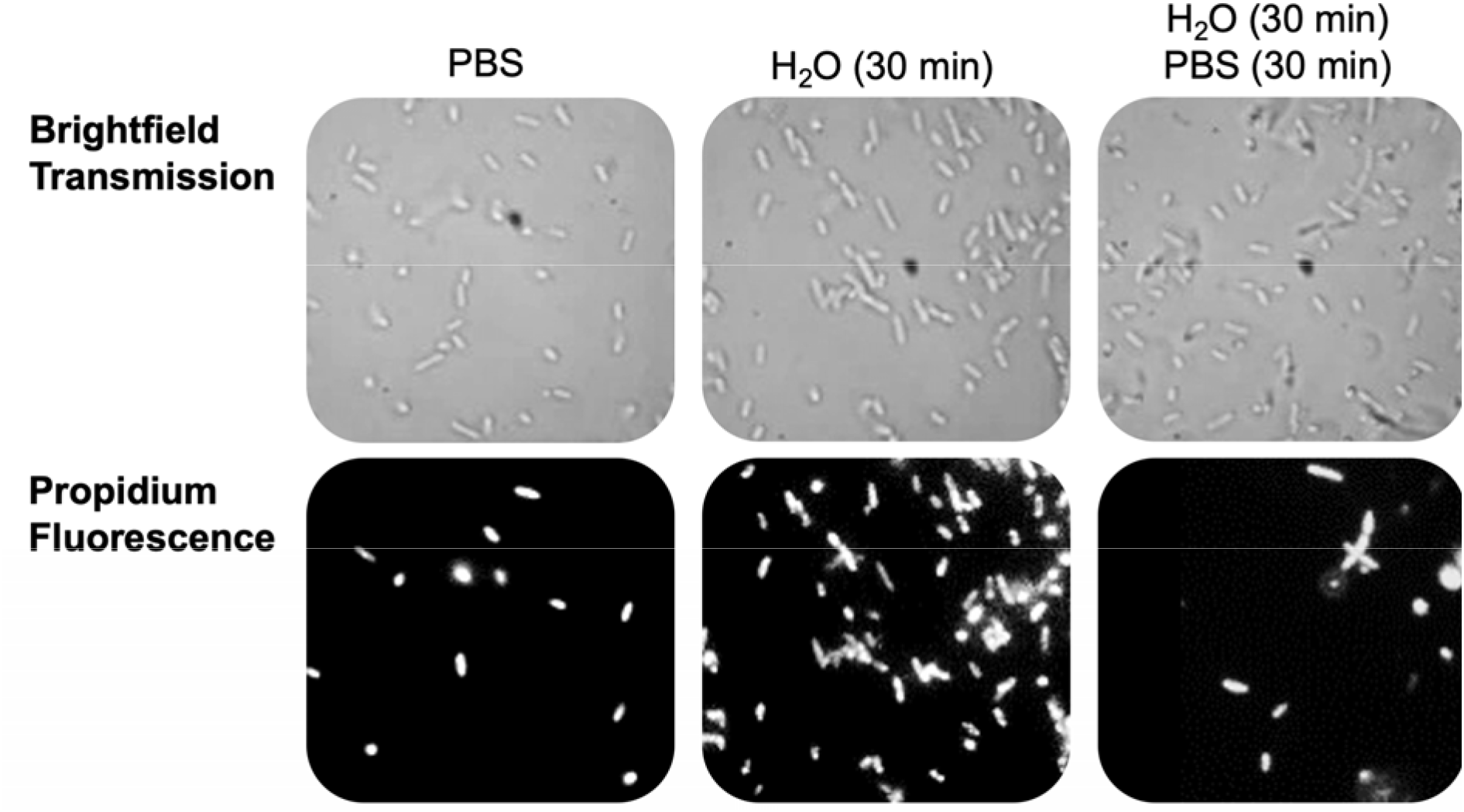
Measured brightfield transmission (top row) and propidium iodide fluorescence (bottom row) images of E. coli under the osmolarity conditions described in **Figure 1**.

## DISCUSSION

In this study, we assessed the influence of hypo-osmotic shock (osmotic downshock) on the structural integrity of *E. coli* membranes using time-resolved SHS. Our study focused on three distinct experimental scenarios, namely: 1) cells exposed to a transient osmotic downshock, for which the MS channels briefly opened then rapidly closed following re-equilibration of osmolarity across the CM, 2) cells exposed to persistent osmotic downshock, for which the MS channels were forcibly held open, and 3) cells exposed to a prolonged osmotic downshock followed by a cessation of shock, for which the MS channels were held open for 30 minutes and then allowed to close.

The state of the MS channels was controlled by the difference in ion concentrations across the CM, which depended on the composition of the suspension medium. When the bacteria were suspended in PBS, the MS channels are expected to remain closed. Conversely, when the bacteria were suspended in distilled deionized water, osmotic pressure created by the ion concentration differentials forced the MS channels to remain open. Likewise, a third scenario was also created, in which cells originally suspended in distilled water (open MS channels) were re-suspended in PBS in order to close their MS channels.

Our lab has previously demonstrated that MG cations freely diffuse across the bacterial CM at a rate two orders of magnitude slower compared to the rate at which they cross the Omp channels in the OM (i.e., k_CM_ = 2×10^-4^ s^-1^ vs. k_Omp_ = 7×10^-2^ s^-1^, **Table 1**).^45,49^ Provided they can be activated, membrane embedded protein channels offer a more efficient means for molecular ions to cross the CM. For example, gating of MS channels not only allows the release of cytoplasmic solutes but also transiently increases the permeability of the CM to external ions.^75^ With an open pore diameter of 1.8 to 3 nm, the MS channels of *E. coli* are more than sufficiently large to facilitate the passive diffusion of a 1 nm wide MG cation. In the SHS measurements presented here, it was observed that by forcing the MS channels to adopt an open state configuration, MG cations were able to cross the CM at a rate at least as fast as the Omp transport rate, ca. k_ms_ ≥ 7×10^-2^ s^-1^. It is noted here that this rate represents a lower bound as k_ms_ transport rates faster than k_Omp_ yield indistinguishable SHS results. Nevertheless, given that both the MS channels and the Omp channels permit non-selective transport of neutral and ionic species, only limited by the width of the open pore diameter, it is reasonable that they would have comparable transport rates. Our results are consistent with previous reports that MS channels in *E. coli* can function as a conduit for ion entry (Ca^2+^, H^+^) and not just osmolyte efflux.^18^

Further, the state of the MS channels was shown to be reversible. The enhanced transport of MG cations observed for open MS channels was subsequently reduced when the MS channels were closed. However, comparison of the SHS traces of the PBS suspended cells (closed MS channels) vs. the cells suspended in water then re-suspended in PBS (open then closed MS channels) suggests that the cells have not yet fully recovered from the prolonged period of osmotic shock. Specifically, as depicted in **Figure 1**, the magnitude and the subsequent decay rate of the CM transport peak are changed following relaxation of persistent shock. While the MS channels quickly close once the applied force is removed, it is likely that the corresponding changes to the cell volume and (likely) lipid packing of the membrane require a substantially longer period to relax to the pre-shock state.^72^

Similar transport behavior was observed for the well-known viability stain, Pro, a CM impermeant molecular cation. In healthy viable cells, Pro is unable to enter the cytosol. In dead or membrane damaged cells, Pro can cross the CM and intercalate with DNA, giving rise to a substantial increase in Pro’s fluorescence intensity. Based upon this simple mechanism, Pro fluorescence imaging can be used to quickly assess dead (fluorescing red) vs. living cells (non-fluorescing). Similar to MG cations, Pro cations are small enough to pass through open MS channels. It was therefore speculated that the general outcome of a Pro fluorescence viability assay could be controlled by modulating whether the MS channels were open. Indeed, as depicted in **Figure 3**, cells suspended in distilled water (open MS channels) fluoresced red, indicating that Pro was able to cross the CM and intercalate with DNA in the cytosol. However, when the cells were then resuspended in PBS (MS channels closed), the number of fluorescing cells was reduced back down to the background level of ca. 20%. Similar to the MG cation SHS kinetics experiments, here too we find that it is possible to significantly increase the transport rate of molecular ions across the CM simply by controlling whether the MS channels are open.

In what follows, we compare and contrast the utility of using patch-clamp (the current gold standard) vs. time-resolved SHS for quantitative experimental observation of MS channel activity. Likewise, we examine the potential for time-resolved SHS to distinguish between MS channels opened due to an applied osmotic shock vs. interaction with a chemical activator.

### A. Patch Clamp vs SHS

The patch-clamp method for monitoring molecular transport across MS channels.^1,4,32–34,38–42^ has a significant limitation in that it is invasive and cannot be applied to the inner membranes of multi-membrane cells, such as Gram-negative bacteria. As demonstrated herein, time-resolved SHS provides a unique ability to selectively monitor molecular transport across open MS channels in suspensions of living bacteria. Further, unlike patch-clamp, time-resolved SHS is not limited to observation of charged molecules. Provided the SHG-active probe molecule can cross the open MS channel (i.e., it is smaller than the pore diameter of the channel of interest), SHS can quantify transport of both neutral molecules and ions across MS channels in suspensions of living cells.

Nevertheless, it is important to acknowledge some of the limitations of SHS (compared to patch-clamp) for monitoring the activity of MS channels. For example, patch-clamp experiments are sensitive to the transport of a single ion across a channel. Each ion crossing event registers as a distinct measured response. Conversely, SHS relies upon the cumulative response of an ensemble of molecules or ions to cross the channel. This stems from the fact that SHS kinetics are not directly sensitive to individual channel crossing events but instead to the sequential adsorption onto the opposing sides of the membrane bilayer containing the channel. Further, in the current demonstration, while it is clear that some MS channels have been activated, the specific identification of these channels (e.g., MscL, MscS) remains uncertain. For instance, considering the distinct pore geometries of MscL and MscS channels, with diameters of 3 nm and 1.8 nm respectively, it is probable that the 1 nm wide MG cations were able to traverse both of these MS channels (as well as the various other MS channels contained in *E. coli*). This underscores the current general lack of specificity associated with this approach. Admittedly, this lack of specificity exists also for patch clamp experiments (i.e., if two different channels where present their behavior would be indistinguishable). However, this issue can be circumvented in patch clamp experiments by selectively isolating the channel of interest. In order to selectively investigate the activity of a specific MS channel in a living cell, enhancements to this methodology could involve a series of carefully chosen bacterial knockout strains (similar to what is already done in some patch-clamp studies). Alternatively, the approach could instead employ SHG-active probe molecules with varying sizes. For example, these probes should be large enough to hinder traversal through the approximately 1.8 nm wide pore of an MscS channel, yet small enough to traverse the approximately 3 nm wide pore of an MscL channel. Likely, some combination of selective knockout strains and probe molecules of distinct sizes would provide the most control in designing such experiments.

### B. MS Channel Open-State Geometry: Osmotic Shock vs. Chemically Activation

Chemical activation of MS channels is currently a topic of active research.^76–79^ The absence of homologues of MscL in the human genome presents an innovative opportunity for antimicrobial targeting.^80^ This approach holds promise for the development of novel antimicrobial strategies, as it offers a viable means for facilitating the passage of otherwise impermeable compounds across the bacterial cell membrane. Note that for an antimicrobial agent to effectively exert its influence, it must have access to the specific biological compartment where it can act. For many antimicrobials, this necessitates traversing the cell membrane and interacting with cytosolic components in order to target metabolomic, replicational, transcriptional, or most significantly, translational pathways.^81^ The rate at which such antimicrobials traverse the cell membrane and accumulate a lethal concentration is a key determinant of the minimum inhibitory concentration (MIC).

In the 1990s, Martinac and colleagues^29,82^ reported that certain amphipathic compounds can activate MS channels in *E. coli*. This discovery was the first identification of MS channel activators and supported the hypothesis that the local population of lipids surrounding the protein complex provide the mechanical gating force needed to open these channels.^3^ In the 2000s, parabens (i.e., alkyl esters of p-hydroxybenzoic acid) were identified as molecules that perturb membrane lateral pressure, activating both MscS^83^ and MscL channels,^78^ suggesting that the antimicrobial action of parabens involves interacting with these channels. Similarly, ‘Compound 10’,^79^ was shown to be a potent antibiotic against drug-resistant Gram-positive bacteria with limited activity against Gram-negative species, presumably due to the presence of the lipopolysaccharide-containing outer membrane. Another noteworthy example is dihydrostreptomycin (DHS), which primarily interferes with protein synthesis in the cytosol.^84–86^ Its effectiveness is reliant on the expression of MscL channels.^76,77^ Dubin *et al*. observed that treatment with DHS resulted in an outward flux of K^+^ prior to any decrease in cell viability, suggesting that the opening of MS channels or loss of cell membrane integrity may be responsible for this phenomenon.^76,87^

The exploration of synergistic effects between MS channel activators and specific antimicrobials holds significant promise within the field of antimicrobial drug discovery. By including an MS channel activator in a combination therapy, it is conceivable that an antimicrobial agent, which would otherwise be unable to penetrate the cell membrane, could gain entry to the bacterial cytosol. This mechanism has the potential to significantly enhance the efficacy of a membrane-impermeable antimicrobial or even substantially reduce the MIC of a compound with weak membrane permeability. Polyvalent ions, on the other hand are reported to be MS channel blockers by affecting on the lateral (in-plane) packing of phospholipid.^88–96^ Our study suggests that SHS could be a useful experimental paradigm for *in vivo* study of channel activators, which is of particular interest in antimicrobial drug discovery.

Finally, SHS could also be used to provide insight for understanding the general structures of MS channels. For instance, a current open question relates to the comparative geometry of MS channels under the influence of osmotic shock vs. chemical activation. Specifically, does chemical activation of MS channels result in a fully open channel (as occurs during osmotic shock) or does local binding of the channel activator simply perturb the configuration of the protein such that these channels become permeable to otherwise impermeant compounds? Time-resolved SHS could be used to differentiate such scenarios. As distinct from the rapid transport of MG cations across an osmotic shock opened MS channel (i.e., exhibiting a two orders of magnitude enhancement in the transport rate), a chemically activated MS channel may not be fully open and therefore may not result in a dramatic enhancement in the MG transport rate. This is conceptually similar to what we recently observed for MG cations crossing the indole-activated mtr channels in the CM of *P. aeruginosa*.^47^ Specifically, in the presence of extra-cellular indole, the transport rate of MG cations across the CM was effectively doubled from 6.5×10^-4^ s^-1^ to 10.6×10^-4^ s^-1^. Importantly, as distinct from the case of osmotic shock activated MS channels (**Figure 1**), in which the CM transport peak is completely lost; the CM transport peak remains observable following only a two-fold enhancement of the transport rate. This behavior can be clearly observed in our simulated SHS responses plotted in **Figure 2**. Specifically, the red curve depicts the scenario in which the MS channels are not contributing to transport across the CM and MG is simply passively diffusing across the CM at a rate of 2×10^-4^ s^-1^. Conversely, the simulation depicted by the orange curve corresponds to transport across the CM at twice the passive diffusion rate of 4×10^-4^ s^-1^ (in addition to the concurrent passive diffusion rate of 2×10^-4^ s^-1^). Consequently, whereas a completely open MS channel results in complete loss of the CM transport peak, chemically activated MS channels may result in more subtle perturbations in the transport kinetic curves.

## CONCLUSIONS

As shown previously, time-resolved SHS can be used to quantitatively monitor molecular adsorption and transport across membranes in living cells. Using this approach, we now demonstrate that by carefully controlling the composition of the suspension media, SHS can be used to observe the action of MS channels in response to applied osmotic pressure. Specifically, we created three distinct sample scenarios: 1) transient osmotic shock, for which the MS channels were closed; 2) persistent osmotic shock, for which the MS channels were forcibly held open; and 3) persistent shock followed by cessation of shock, in which the MS channels were held open for a prolonged duration and then allowed to close. In this study, the second harmonic active molecular ion, malachite green, was used as the probe of transport across the *E. coli* cytoplasmic membrane.

When the MS channels were closed (scenarios one and three), slow transport across the bacterial CM was observed, characteristic of gradual diffusion across the hydrophobic interior of the CM. Conversely, when the MS channels were held open (scenario two), rapid transport across the CM was observed (i.e., at least two orders of magnitude faster compared to passive diffusion). These observations were corroborated by complementary fluorescence experiments using the Pro cation.

Unlike the patch-clamp experiment, SHS holds the promise of being able to experimentally observe the action of MS channels in structurally intact / living cells. Further, SHS can likely be used to understand structural changes to MS channels following chemical activation, as compared to shock-induced activation of MS channels.

## AUTHOR INFORMATION

### Notes

The authors declare no competing financial interest.

## ACKNOWLEDGEMENTS

This work was supported by the National Science Foundation grant CHE-2105387. We wish to thank Charles D. Cox (Victor Chang Cardiac Research Institute) for helpful discussions which inspired this work.

## ABBREVIATIONS

MS: mechanosensitive
MscL: mechanosensitive channel of large conductance
MscS: mechanosensitive channel of small conductance
SHS: second-harmonic light scattering
MG: malachite green
OM: outer membrane
CM: cytoplasmic membrane
PM: peptidoglycan mesh
*E. coli*: *Escherichia coli*.

